# Additive Dose Response Models: Explicit Formulations and the Loewe Additivity Consistency Condition

**DOI:** 10.1101/161059

**Authors:** Simone Lederer, Tjeerd M. H. Dijkstra, Tom Heskes

## Abstract

High-throughput techniques allow for massive screening of drug combinations. To find combinations that exhibit an interaction effect, one filters for promising compound combinations by comparing to a response without interaction. A common principle for no interaction is Loewe Additivity which is based on the assumption that no compound interacts with itself and that doses of both compounds for a given effect are equivalent. For the model to be consistent, the doses of both compounds have to be proportional. We call this restriction the Loewe Additivity Consistency Condition (LACC). We derive explicit and implicit null reference models from the Loewe Additivity principle that are equivalent when the LACC holds. Of these two formulations, the implicit formulation is the known General Isobole Equation [1], whereas the explicit one is the novel contribution. The LACC is violated in a significant number of cases. In this scenario the models make different predictions. We analyze two data sets of drug screening that are non-interactive [2, 3] and show that the LACC is mostly violated and Loewe Additivity not defined. Further, we compare the measurements of the non-interactive cases of both data sets to the theoretical null reference models in terms of bias and mean squared error. We demonstrate that the explicit formulation of the null reference model leads to smaller mean squared errors than the implicit one and is much faster to compute.

## I. Introduction

In mixture toxicology and compound interaction modeling one is interested in synergistic or antagonistic effects between biological compounds. When combining two or more compounds, their combined effect can be much larger than the individual effects. Such a so-called synergistic effect allows for administration of lower doses to reach the same effect. This has applications in many areas such as chemotherapy [4].

The basic understanding of synergy is any effect greater than the expected effect with no interaction assumed. This expected effect without interaction is specified with a so-called null reference model. Therefore, synergy depends highly on such a reference model of a non-interactive scenario. The central problem of defining such null reference models is the prediction of a response surface from the conditional responses. Conditional responses are the responses to a single compound, that is, conditional on the concentration of the other compound being zero.

Throughout the last century, several models for the null reference and methods to measure the deviance from these have been proposed. An extensive overview is given by Greco *et al.* [5] and recent reviews are given by Geary [6] and Foucquier and Guedj [7]. One of the most famous null reference models is the General Isobole Equation, which was introduced by Loewe [1], and is based on the so-called Loewe Additivity principle. Several other models have been introduced such as Bliss Independence [8], Chou and Talalay’s method [9], which concentrate on the null reference model locally, and the ZIP model [2]. Despite the variety of null reference models, there is no agreement on a best model or a best practice on how specifically synergy is detected. However, Loewe Additivity enjoys a wide reputation because of its principle of the sham combination. This principle rests on the idea that a compound combined with itself should yield no interaction effect.

As the first experiments for the assessment of synergy were conducted in vivo, one used to administer varying doses of compounds, that is, the unit of compound per kilogram of biological system under investigation. That historical term still remains in the research area of synergy and often the term dose is used to actually refer to concentrations, the number of molecules per unit volume. Throughout this study, we refer to the measured effect of a compound combination both as response and effect. In the literature this measured effect is also referred to as the phenotypic effect or as cell survival of disease agents or cancer cell lines. Measurements taken for only one compound, here referred to as the conditional responses, are also called mono-therapeutic [10] or single compound, but we prefer a more statistical terminology. We refer to the measurements of one cell line exposed to all combinations of the two compounds as a record, but in other literature it is referred to as response matrix [2, 11].

Loewe Additivity is a phenomenological description, not a mechanistic one that is aiming to explain underlying mechanisms. A way to root Loewe Additivity in such mechanistic terms is undertaken by Baeder et al. [12]. Further, we do not take temporal effects into consideration but work uniquely in the concentration space. While temporal considerations are important, in most high-throughput studies the effect is measured after a fixed period when transient responses have died out, but before effects like cell division set in.

We first give a short introduction to conditional dose response curves in section II, to then describe the most common null reference principle, Loewe Additivity, in section III. We study Loewe Additivity’s consistency condition and its consequences. Further, we introduce an explicit null reference model derived from the Loewe Additivity principle, which describes the same null reference model as the General Isobole Equation, when the Loewe Additivity consistency condition is met. As this consistency condition is often violated by experimental data [6, 13] we investigate in section IV the consequences of these violations and provide solutions to the arising issues, which we evaluate in section V.

## II. Conditional Dose Response Curves

A common approach for modeling monotonic doseresponse curves *f*_*j*_ with *j* ∈{1, 2} is the Hill curve [14], also referred to as sigmoid function. The Hill model is, due to its good fit to many sources of data, the most widely applied model for fitting compound responses [15]. It has a sigmoidal shape with little change for small doses but with a rapid decline in response once a certain threshold is met. For even larger doses the effect asymptotes to a constant maximal effect. Two exemplary Hill curves are depicted in Fig. 1. There are several parameterizations of the Hill curve. We use the following throughout this study to fit conditional responses:

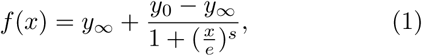

where *y*_0_ is the response at zero dose and *y*_*∞*_ the maximal response of the cells to the compound, *e* the dose concentration reaching half of the maximal response and *s* the steepness of the curve. Eq. 1 is equivalent to the parametrization used in the drc package [16], the so-called four parameter log-logistic model. By our definition of the Hill curve, a positive *s* leads to a descending Hill curve.

**Figure 1:**
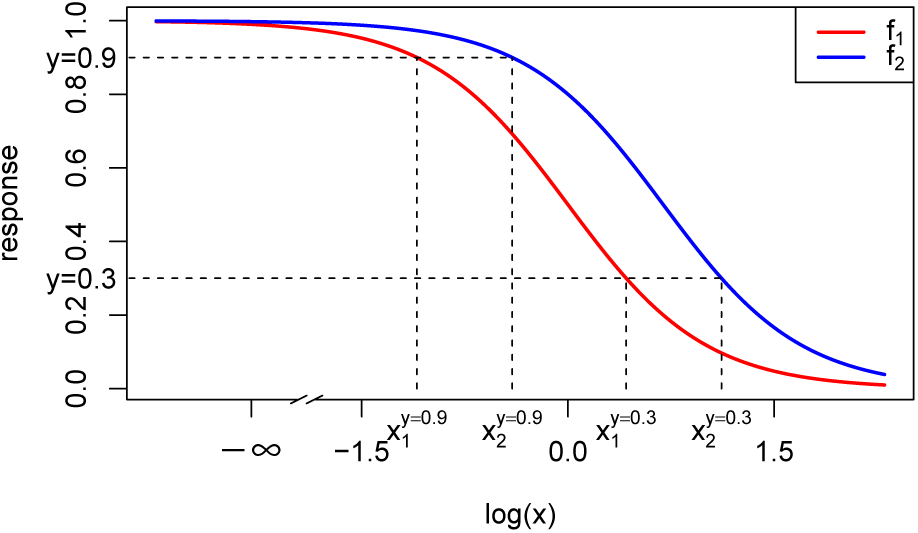
Dose-response curves (red and blue) as Hill curves (Eq. 1). For the exemplary responses of 0.3 and 0.9 the different doses x_1_ and x_2_ reaching that effect are shown (dashed lines). The doseresponse curves differ only in EC50 with e_1_ = 2 and e_2_ = 1. Values of the other parameters are y_0_ = 1, y∞ = 0 and s = 2. To highlight the sigmoidal shape of a Hill curve in log-space, the logarithmic concentration space is depicted.

## III. Null Reference Models

To quantify the degree of synergy between two compounds, their measured combination effect is compared to an expected effect assuming no interaction. The larger the deviance to such a null reference model, the larger the interaction effect. When combining two compounds, one typically measures also the response of each compound in the absence of the other. Knowing these two conditional responses of both compounds, the aim of a null reference model is to define the surface spanned between these conditional responses, assuming no interaction between them.

Throughout the extensive research that was conducted in the field of synergy over the last century, several null reference principles were introduced, but only two survived the critics [5]: Loewe Additivity [1] and Bliss Independence [8]. The first assumes that one compound can be substituted for another (with a certain ratio dependent on each compound’s efficacy). As an interpretation one can say that according to Loewe Additivity the compounds have the same mechanism of action (act on the same pathway). In Bliss Independence, one assumes that the responses can be added as the compounds have a different mechanism of action (act independently). Throughout this paper we will exclusively focus on Loewe Additivity as it was shown that Loewe Additivity predicts synergy better than Bliss Independence [3].

The Loewe Additivity principle is based on an experiment with a sham combination, that is, combining a compound with itself. A compound does not interact with itself and yields therefore the same effect as if only one compound was administered with the sum of the doses. Combining two compounds, it is implied that these “act similarly, presumably at the same site of action, differing only in potency”, where potency is meant to be the effect [5, p.344]. To clarify this idea, consider an effect of *y* = 0.3, as depicted in Fig. 1 with the lower dashed horizontal line. An effect of *y* = 0.3 denotes therefore the survival of 30% of the cell culture. We denote 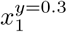 to be the dose of compound 1 that reaches an effect of *y* = 0.3 and 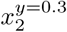 the dose of compound 2 to reach an effect of *y* = 0.3. In Fig. 1, these are depicted on the *x*-axis where the two most right vertical dashed lines intersect, which are descending from the *y* = 0.3 effect that is reached by the red and blue curve, respectively.

When visualizing the null response surface that is spanned between the two Hill curves the doses of the two compounds are mapped onto the *x*- and *y*-axis of a Cartesian coordinate system and the response surface can be represented in form of a contour plot, yielding a 2D representation, see Fig. 2. Also here, one can indicate the doses 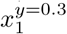 on the *x*- and 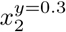 on the *y*-axis. Loewe argues that any compound combination that lies on the straight line drawn between these two points on the axes should also yield an effect of *y* = 0.3. Mathematically, this corresponds to any set of dose pairs (*x*_1_*, x*_2_) of the two compounds, which, divided each by the doses that reach an effect of *y* = 0.3 individually, sum up to one:

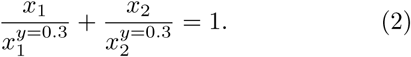

**Figure 2:**
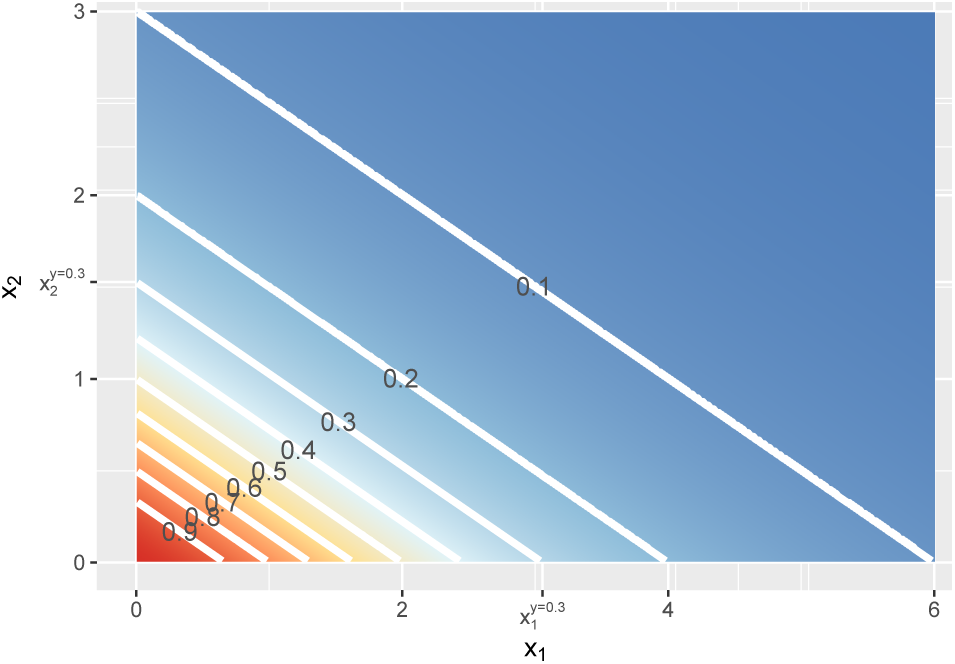
Contour lines of the response surface from Eq. 5, 8 or 9 with x_1_ on the x-axis and x_2_ on the y-axis at linear concentrations.

This idea can be generalized from *y* = 0.3 to any effect *y*, which follows a continuous response curve *f*_*j*_(*x*_*j*_) with *j ∈* {1, 2} for each compound:

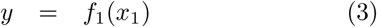

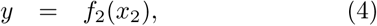

with a common response for the control dose *f*_1_(0) = *f*_2_(0) and functions *f*_1_ and *f*_2_ being monotonic (either increasing or decreasing). If doses 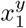 and 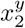 individually each reach the same effect *y*, the dose 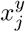 can be written as the inverse of the response curve, namely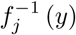. This yields the so-called General Isobole Equation [1, p.179] [17], in the following referred to as *f*_GI_ (*x*_1_*, x*_2_):

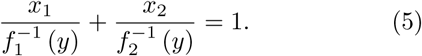

This defines a response surface as depicted in Fig. 2 as a contour plot. Different isoboles, also known as iso-effect curves for response surfaces, are depicted for different effects *y* and labeled in the plot.

For specific forms of *f*, the General Isobole Equation model *f*_GI_ (*x*_1_*, x*_2_) can be computed analytically. In general, and in particular if *f* takes the form of a Hill curve, the definition of *f*_GI_ (*x*_1_*, x*_2_) is implicit and numerical computations are needed to solve Eq. 5. More details on how to solve Eq. 5 numerically are presented in Appendix E.

Following the main idea of Loewe Additivity presented in [1], we formalize a trade-off between the two compounds, meaning that we can exchange a given dose of compound 2 (*x*_2_) for a dose of compound 1 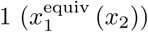 with the equivalent effect. Vice versa, we can exchange a given dose of compound 1 (*x*_1_) for an effect equivalent dose of compound 2 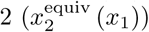

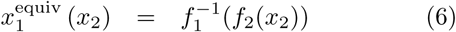

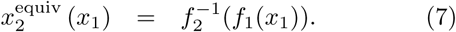

The construction of response equivalent doses is illustrated in Fig. 1 for response levels *y* = 0.9 and *y* = 0.3. With these equivalent doses one can construct two response surfaces. To do so, we add to the concentration *x*_1_ of compound 1 its equivalent dose 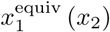 and the same mutatis mutandis for *x*_2_ and compute their effect:

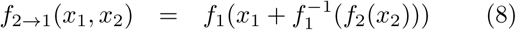

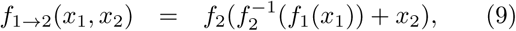

where effect *f* _*j→k*_ stands for converting the dose of compound *j* into the equivalent dose of compound *k*. The response surfaces *f*_2*→*1_(*x*_1_*, x*_2_) and *f*_1*→*2_(*x*_1_*, x*_2_) are plotted in Fig. 2 as a contour plot, spanning a surface between the first and second compound, that are depicted on the *x*- and *y*-axis, respectively. Note that the response surfaces from Eq. 8 and Eq. 9 are identical to the surface from Eq. 5. We will expand on this finding later.

The two formulas in Eq. 8 and Eq. 9 must be equivalent as both span the surface by expressing the concentration of one compound in terms of the other compound yielding the same effect. Thus we introduce the Loewe Additivity Consistency Condition, abbreviated as LACC, namely that both equations, Eq. 8 and Eq. 9, must be equivalent:

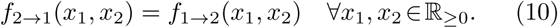

To our knowledge, we are the first to explicitly state this consistency condition in a general mathematical form. Surprisingly and often ignored in literature (e.g. [2, 5]), the two formulas in Eq. 8 and Eq. 9 do not always yield the same surface which leads to violations of the LACC. In Theorem 1 we show that, in order for these two equations to be equal, the dose and its effect equivalent dose have to be proportional to each other:

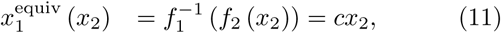

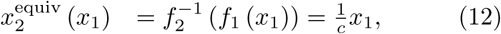

for a constant *c >* 0. The violation of the consistency condition in Eq. 10 by violating the condition in Eq. 11 and 12 has been commented upon before by Tallarida and Geary [6, 13], but no general proof has been proposed.

### Theorem 1.

*If and only if a dose and its equivalent are proportional to each other (Eq. 11 and 12) the Loewe Additivity Consistency Condition in Eq. 10 holds.*

The proof can be found in Appendix A. Note that in the proof, the important implicit assumption is made, that both response curves yield the same maximal effect *y*_*∞,*1_ = *y*_*∞.*2_. In further discussion of the LACC we assume this equality. In section IV, we investigate the consequences if the condition is not met.

Therefore, the principle of Loewe Additivity only makes a consistent prediction for the response to a mixture of compounds when the trade-off between the compounds is linear. Put another way, the LACC is only fulfilled when the dose-response curves are shifted copies of each other on the logarithmic dose axis since 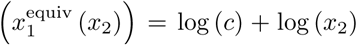 This shift is depicted in Fig. 1. Two Hill curves are depicted, where one is shifted horizontally relative to the other. The vertical dashed lines indicate the different doses that reach the same effect, represented by the horizontal dashed lines at the exemplary effect levels of *y* = 0.3 and *y* = 0.9. The comparison of these two effects shows that equivalent doses are shifted equidistantly, independent of the effect they reach.

As a corollary to Theorem 1, we show that isoboles are linear and parallel when the LACC holds.

### Corollary 1.

*If the Loewe Additivity Consistency Condition in Eq. 10 holds,* (1) *f*_*GI*_ (*x*_1_*, x*_2_) = *f*_2*→*1_ (*x*_1_*, x*_2_) = *f*_1*→*2_ (*x*_1_*, x*_2_) *and* (2) *the isoboles are parallel.*

The proof can be found in Appendix B. Applying the LACC in Eq. 11 with *f*_1_ and *f*_2_ taking the form of two Hill curves, the following restriction on Hill curves guarantees the LACC (Eq. 10) to hold:

### Corollary 2.

*If the Loewe Additivity Consistency Condition in Eq. 10 holds with f*_1_ *and f*_2_ *taking the form of two Hill curves, then the slopes s and effect ranges y*_0_ *and y*_*∞*_ *of the Hill curves must be the same. Further, the proportionality factor c takes the form of a fraction of the EC50 value e of the drug to be expressed in terms of the other divided by the EC50 value of this other drug, resulting in* 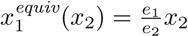

See Appendix C for the proof.

In summary, we have now derived three possible null reference models for Loewe Additivity. These are *f*_2*→*1_(*x*_1_*, x*_2_) (Eq. 8), *f*_1*→*2_(*x*_1_*, x*_2_) (Eq. 9) and *f*_GI_ (*x*_1_*, x*_2_) (Eq. 5) which are all equivalent under the LACC.

Note that the first two models, *f*_2*→*1_(*x*_1_*, x*_2_) (Eq. 8) and *f*_1*→*2_(*x*_1_*, x*_2_) (Eq. 9) are explicit formulations of the response surface and *f*_GI_ (*x*_1_*, x*_2_) (Eq. 5) is implicit. Thus, there is an arbitrariness to Loewe Additivity which allows the construction of other null reference models.

## IV. Violation of the Loewe Additivity Consistency Condition

In this section we investigate the consequences of violations of the LACC. As mentioned before and commented by Tallarida [13] and Geary [6], the conditional response curves of experimental data are often not parallel and therefore, the LACC in Eq. 10 is often violated. Here, we investigate what different violations of the LACC imply for the null reference models.

In Fig. 3, two Hill curves are depicted in three scenarios where the LACC is violated: two different slopes *s* or two different maximal effects *y*_*∞*_ or both. It becomes immediately clear that in all scenarios there is no proportional relationship between the two curves. In Fig. 4 the three null reference models *f*_GI_ (*x*_1_*, x*_2_), *f*_2*→*1_ (*x*_1_*, x*_2_) and *f*_1*→*2_ (*x*_1_*, x*_2_) are depicted with the conditional responses following the Hill curves as depicted in Fig. 3. The *f*_GI_ (*x*_1_*, x*_2_) model is depicted in Fig. 4a, while the two explicit models, *f*_2*→*1_(*x*_1_*, x*_2_) and *f*_1*→*2_(*x*_1_*, x*_2_) from Eq. 8 and Eq. 9, are depicted in Fig. 4b and Fig. 4c, respectively. In each column, the three cases of LACC violation are depicted.

**Figure 3:**
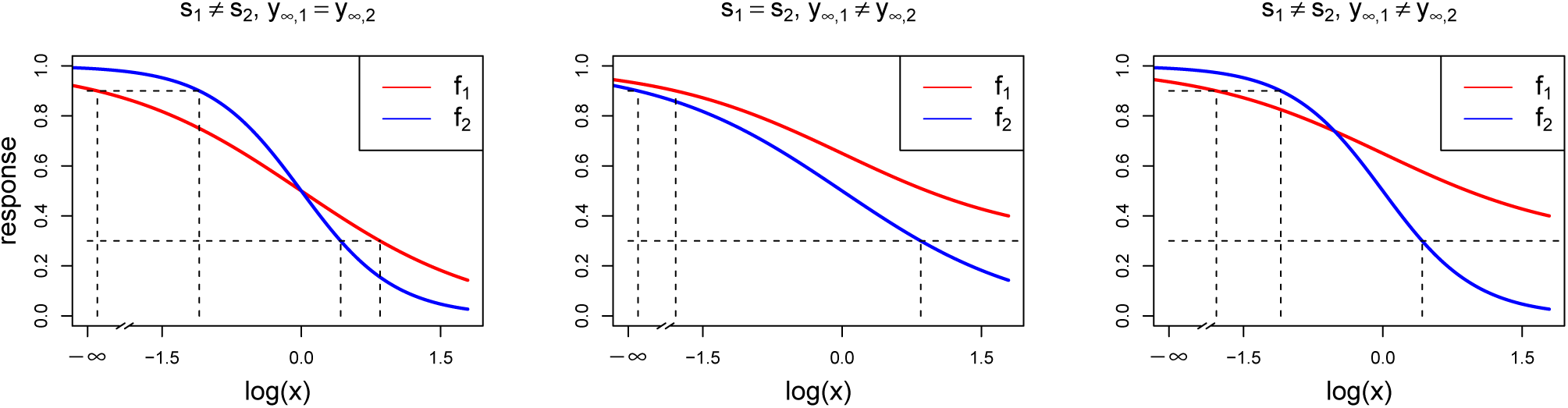
Dose-response curves (red and blue) with different parameter settings that all violate the LACC: s_1_≠s_2_ (left) with s_1_ = 1 and s_2_ = 2, y∞,_1_ ≠y∞,_2_ (center) with y∞,_1_ = 0.3 and y∞,_2_ = 0, and (right) with s_1_ ≠s_2_ and y∞,_1_≠y∞,_2_ with the same settings as above. Additionally, all other parameters that are chosen to be equal for both Hill curves take the values: s = 1, y_0_ = 1, y∞ = 0 and e = 1.

**Figure 4:**
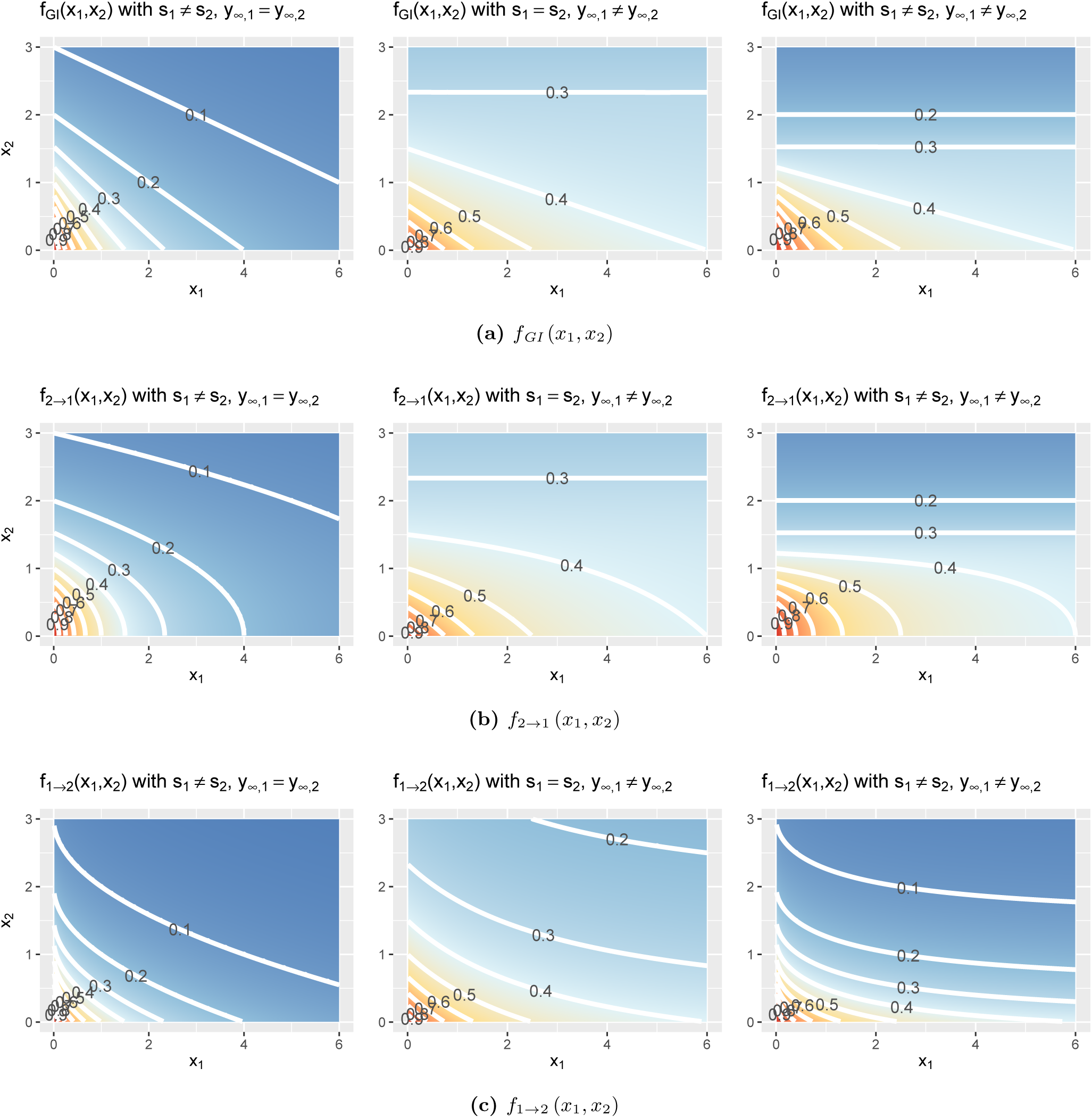
Violation of the LACC with either different slope parameters s_1_ ≠s_2_ (left), or different maximal effects y∞ (middle) or both (right) for the fGI (x_1_, x_2_) (a), f_2_→_1_ (x_1_, x_2_) (b) or f_1_→_2_ (x_1_, x_2_) (c) model.

Let us first investigate in detail the first case of violation, assuming different slope parameters for the conditional responses, as depicted in the left column of Fig. 4. The *f*_GI_ (*x*_1_*, x*_2_) displays straight isoboles, which are not parallel as they are in Fig. 2. The straightness is due to Berenbaum’s definition of the General Isobole Equation and becomes obvious by inspection of Eq. 5, which is symmetric in the fractional terms. This is one of the reasons why this model has been popular. The two explicit models in the left panel of Fig. 4b and Fig. 4c display a concave or convex curvature to the point of zero dose concentration.

Taking into consideration the common violation of the LACC, as we show in section V, and the resulting difference in response surfaces spanned by the three null reference models, as depicted in the left column of Fig. 4, we additionally define a new null reference model, that is equivalent to the General Isobole Equation model under the LACC, as a linear combination of the two explicit formulations. Further, even under the violation of the LACC in case of different slopes this combination gives almost straight isoboles in the response surface: Since the General Isobole Equation is clearly symmetric, we take a weighted mean of the two explicit formulations *f*_2*→*1_ (*x*_1_*, x*_2_) and *f*_1*→*2_ (*x*_1_*, x*_2_) given in Eq. 8 and Eq. 9:

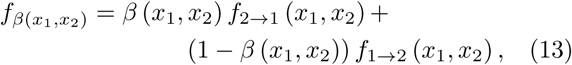

with *β* (*x*_1_*, x*_2_) *∈* [0, 1].

We show in Appendix D that equal weights are most insensitive to a violation of the LACC by differing slopes:

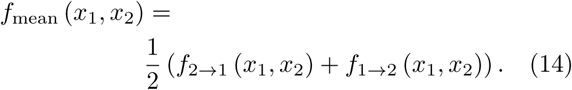

In the following, we will refer to Eq. 14 as the Explicit Mean Equation, or mathematically, as *f*_mean_ (*x*_1_*, x*_2_). A visualization of *f*_mean_ (*x*_1_*, x*_2_) is given in Fig. 2 if the LACC holds and if the LACC is violated the behaviour of *f*_mean_ (*x*_1_*, x*_2_) is shown Fig. 5. For each of the three scenarios of violation of the LACC, as depicted in Fig. 3, *f*_mean_ (*x*_1_*, x*_2_) is shown. The contour lines of the *f*_mean_ (*x*_1_*, x*_2_) are depicted in white and for reference, the contour lines of *f*_GI_ (*x*_1_*, x*_2_) are depicted in grey. In the left panel of Fig. 5, the situation is depicted for different slopes *s* but the same *y*_*∞*_. For this violation of the LACC the model shows nearly linear isoboles, with a slight curvature which is almost linear for *x*_1_ reaching a larger effect than *x*_2_ and convex for *x*_1_ reaching a smaller effect than *x*_2_. The other two cases of violation of the LACC will be discussed below.

**Figure 5:**
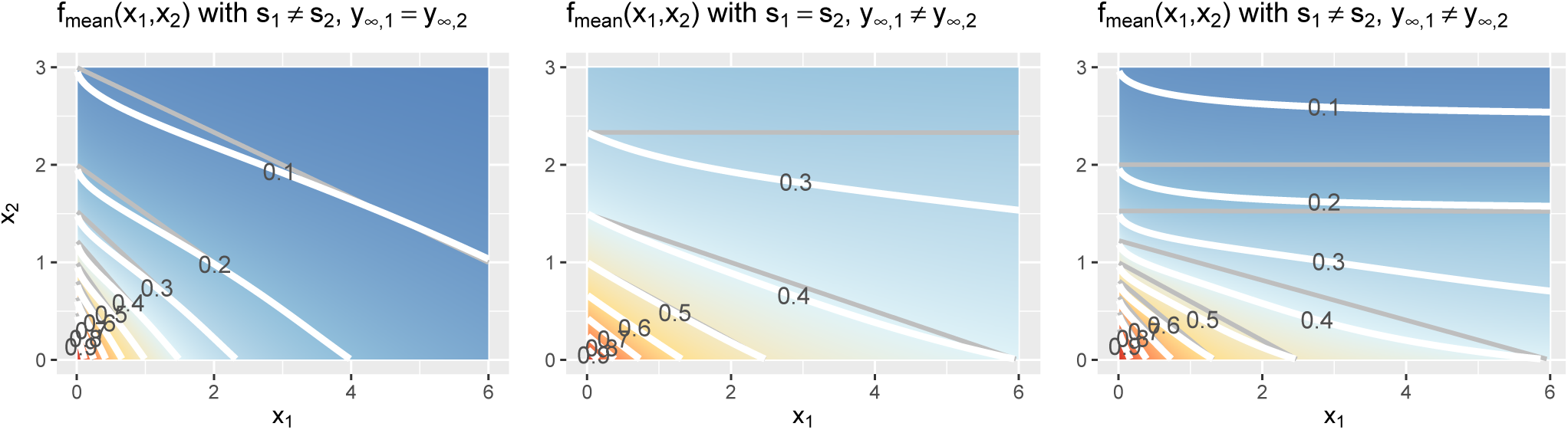
Contour lines of the fmean (x_1_, x_2_) model depicted in white and the corresponding contour lines of fGI (x_1_, x_2_) shown in grey. The three different scenarios, in which the LACC is violated, are shown from left to right: the slopes are different, s_1_ = s_2_, here depicted with s_1_ = 1, s_2_ = 2, y_∞_ = 0, or the maximal effect values differ, y_∞,1_ = y_∞,2,_ here shown with s = 1, y_∞,1_ = 0.3, y_∞,2_ = 0 or both are different, here shown with s_1_ = 1, s_2_ = 2, y_∞,1_ = 0.3, y_∞,2_ = 0. The remaining two parameters of the Hill curve are set equal for all figures to y_0_ = 1 and e = 1.

Above we mentioned that the slopes often differ, and it turns out that the same holds for the maximal effect values. In case one dose reaches an effect that cannot be reached by the other, there is no equivalence relationship between the two doses and therefore Loewe Additivity is not defined. Assume that dose *x*_1_ reaches a stronger effect that cannot be reached by compound 2:

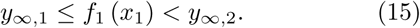

For that effect *y* = *f*_1_ (*x*_1_), the inverse of the Hill curve of compound 2, 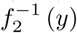 is not defined. For the *f*_GI_ (*x*_1_*, x*_2_) model, di Veroli [10] suggests that one would need an infinite dose of *x*_2_ to reach that effect *y*, resulting in 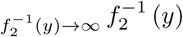 and inserted into the *f*_GI_ (*x*_1_*, x*_2_) model gives:

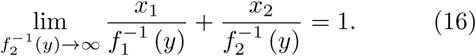

As the second fraction in the sum on the left-hand side vanishes, the expected response of the *f*_GI_ (*x*_1_*, x*_2_) model for that dose combination, is the effect that is reached by compound 1 alone: *y* = *f*_1_ (*x*_1_).

The inverse of the Hill curves of compound 2 is not defined for any effect *y* greater than the maximal effect *y*_*∞,*2_. This has as consequence for the explicit models that *f*_1*→*2_ (*x*_1_*, x*_2_) is not defined. Following a similar line of thought as di Veroli [10]:

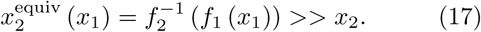

Therefore, within 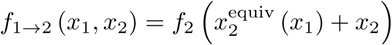 *, x*_2_ becomes negligible. This results in

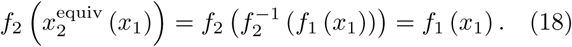

Therefore, we suggest for *f*_mean_ (*x*_1_*, x*_2_) to compute the effect that is reached by the mean of *f*_2*→*1_ (*x*_1_*, x*_2_) and *f*_1_ (*x*_1_):

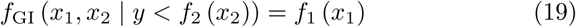

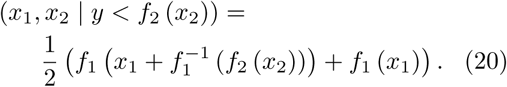

Thus *f*_mean_ (*x*_1_*, x*_2_) predicts a larger effect than the *f*_GI_ (*x*_1_*, x*_2_) model. This solution is depicted in the middle panels of Fig. 4 for the *f*_GI_ (*x*_1_*, x*_2_) model as well as for *f*_2*→*1_ (*x*_1_*, x*_2_) and *f*_1*→*2_ (*x*_1_*, x*_2_), and for the *f*_mean_ (*x*_1_*, x*_2_) model in the middle panel of Fig. 5. The models have all a smaller effect than in the previous scenario, where the slopes differ. All explicit models exhibit nonlinear isoboles. *f*_2*→*1_ and *f*_1*→*2_ exhibit a similar concave and convex curvature behaviour as in the scenario of differing slopes. For *f*_GI_ (*x*_1_*, x*_2_), *f*_2*→*1_ (*x*_1_*, x*_2_) and *f*_mean_ (*x*_1_*, x*_2_), the asymptotic behaviour is depicted.

These figures display a constant horizontal response for *x*_1_, only decreasing for increasing *x*_2_ doses. For the case where both scenarios of a violation are met, namely the violation of LACC and non-definition of Loewe Additivity, we depict the response surfaces of the null models in the right column of Fig. 4 and the right panel of Fig. 5. The surfaces are, as in the first case, more contracted to the origin. The curvature behaviour of the isoboles of the explicit models is again similar to both previous cases and the asymptotic behaviour where one dose reaches a higher effect shows again no change in response in the direction of *x*_1_ doses.

Analogously to the weighted mean, we can take the geometric mean of the two explicit models *f*_2*→*1_ (*x*_1_*, x*_2_) and *f*_1*→*2_ (*x*_1_*, x*_2_). This model together with an analogous analysis to the Explicit Mean Equation model that was presented in this section can be found in Appendix G.

## V. Evaluation

With two data sets we will confirm Geary’s statement about the common violation of the LACC. Furthermore, we compare the different null reference models by considering non-interactive records of two different data sets of compound screenings.

The first data set was created by Mathews Griner et al. [18] and is a cancer compound synergy study. We refer to this data set as the Mathews Griner data. It is composed of 463 different drug-drug-cell combinations on the cancer cell line TMD8 and was published along with many other large compound-drug-cell combination studies on the website https://tripod.nih.gov/matrix-client/. It is a so-called one-to-all experiment, meaning that one compound (in this case ibrutinib) is combined with 463 other compounds. In this highthroughput study all 463 compound combinations are screened in a 6 × 6 matrix design and the effect of the compound combinations is measured as cell viability. The six different concentrations of ibrutinib and the paired compound decrease from 2.5μM and 125μM four times with a four-fold dilution with the sixth dose being zero [2].

In a synergy modeling study, Yadav et al. [2] categorized each record of the Mathews Griner data into three interaction classes after visual inspection of its doseresponse matrix: synergy, no interaction, antagonism. We only use the 252 dose response matrices classified as non-interactive.

The other data set used in this study with a labeling of the records is the anti fungal cell growth experiment on the yeast S.cerevisiae by Cokol et al. [3] and from here on referred to as the Cokol data set. In this study 200 different drug-drug-cell combinations were conducted with 33 different compounds and growth inhibition was measured. An 8 × 8 factorial design is used with doses linearly increasing from 0 up to a dose close to the individually measured maximal effect dose of the compound under investigation.

The categorization of this data set is based on a comparison of the longest arc length of an isobole relative to the expected longest linear isobole in a non-interactive scenario. In more detail, having estimated the response surface of a record, Cokol et al. chose the longest contour line and measure its length and direction (convex or concave). In case of the contour line being convex the record is categorized as synergistic and the arc length of the longest contour line determines the strength of synergy. As the labeling of these records is quantitative, we consider all records to be non-interactive if their absolute value is smaller than 0.8. This decision is based on communication with the authors [3]. This leaves us with 82 records.

We received both categorizations after personal communication with the authors [2, 3]. For the purpose of comparing the null reference models introduced in section III and IV, we consider these two classifications as ground truth, given that no molecular information is available for verification.

The conditional responses are fitted with Hill curves individually, with the constraint to share the *y*_0_ parameter. For the fitting, we make use of the drc package [16]. More detailed information is given in Appendix E. Records with negative slopes or negative EC50 values are excluded, which leaves us with 159 records for the Mathews Griner and 79 for the Cokol data. The main reason for the exclusion of nearly 40% of the records of the Mathews Griner data is the fixed dose range applied to all compounds. Many conditional readouts show barely any response over the entire dose range. The compounds might therefore have no effect at all on the cell line or the dose range is too small to cause any effect.

To support the statement from section IV about the LACC being often violated, we apply a Wilcoxon signedrank test to the Mathews Griner data which combines one compound with a set of other compounds. One tests the null hypothesis that the slopes *s* from the two fitted conditional responses are equal. The other one tests for equality of the maximal effects *y*_*∞*_. Both tests on *s* and *y*_*∞*_ are significant with *p*_*s*_= 6.26 − 10^*-*5^ and *p*_*y∞*_ = 2.2 × 10^*-*16^.

To compare the performance of the models to capture no interaction, we compare *f*_GI_ (*x*_1_*, x*_2_) (Eq. 5) and *f*_mean_ (*x*_1_*, x*_2_) (Eq. 14) by computing the bias and mean squared error between the null reference surfaces that are spanned by the models and the measured response data, excluding the outliers (see Appendix E).

Scatter plots of the bias of both data sets are depicted in Fig. 6. For every record, the bias of *f*_mean_ (*x*_1_*, x*_2_) is depicted on the *x*- and the bias of *f*_GI_ (*x*_1_*, x*_2_) on the *y*-axis. A striking observation from Fig. 6 is that the bias values of *f*_GI_ (*x*_1_*, x*_2_) are always larger than those of *f*_mean_ (*x*_1_*, x*_2_). This holds for both data sets. For positive bias values, this gives a smaller bias for *f*_mean_ (*x*_1_*, x*_2_) and for negative bias values, a smaller bias in absolute terms for *f*_GI_ (*x*_1_*, x*_2_). As the bias is the mean of differences of estimated data points to the measured ones, we look in detail into those differences for each record. For all records of the Mathews Griner and most records for the Cokol data set, we find that the individual error values of each data point are larger for the *f*_GI_ (*x*_1_*, x*_2_) model. We suspect this to be due to the definition of the explicit models, as, by taking into consideration the effect of the other compound as well, the spanned surfaces are steeper decreasing if the LACC is violated. This becomes clear by inspecting the right panel of Fig. 5, for which the contour lines of the *f*_mean_ (*x*_1_*, x*_2_) model are depicted in white with the contour lines of the *f*_GI_ (*x*_1_*, x*_2_) model depicted in gray: due to the white contour being more contracted to the origin than the gray ones, the response surface of the explicit *f*_mean_ (*x*_1_*, x*_2_) model has a steeper decrease than the implicit *f*_GI_ (*x*_1_*, x*_2_) model. This is also in line with the negative bias values, which are larger for the *f*_mean_ (*x*_1_*, x*_2_) model in absolute terms. If the *f*_GI_ (*x*_1_*, x*_2_) model spans a surface below the measured data, the *f*_mean_ (*x*_1_*, x*_2_) model then definitely spans a surface below the *f*_GI_ (*x*_1_*, x*_2_) model and therefore below the measured data.

**Figure 6:**
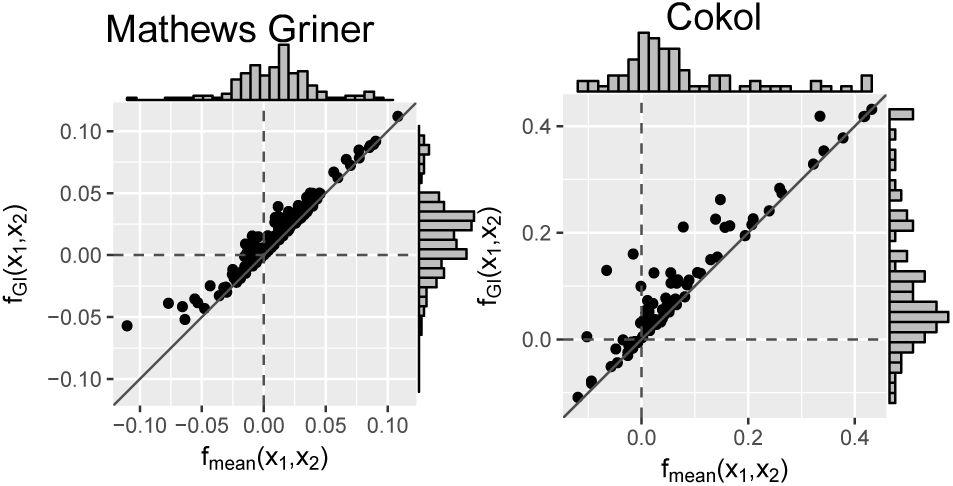
Bias: mean difference between the responses given by the model and the measured responses with the diagonal is depicted. The distribution of the models’ errors is given in histograms plotted on the axes.

Additionally, we compare the mean squared error values to the *f*_GI_ (*x*_1_*, x*_2_) model and the *f*_mean_ (*x*_1_*, x*_2_) model. We do so using the Wilcoxon signed-rank test for paired samples. For the Mathews Griner and the Cokol data, the Wilcoxon signed-rank tests are significant to reject the null hypothesis of an equal mean for the alternative hypothesis that the error values of the *f*_GI_ (*x*_1_*, x*_2_) model are greater. The *p*-value for the errors of both null models on the Mathews Griner data is *p*_Mathews Griner_ = 1.46 − 10^*-*6^ and for the Cokol data is *p* _Cokol_ = 7.68 − 10^*-*5^. Further, in Fig. 7, the errors of both data sets are depicted in two scatter plots, the Mathews Griner data on the left and the Cokol data on the right hand side. The mean squared errors of the *f*_mean_ (*x*_1_*, x*_2_) model are drawn on the *x*-axis and the mean squared errors of the *f*_GI_ (*x*_1_*, x*_2_) model on the *y*-axis. The models are considered to perform equally well if their mean squared error values are equal and therefore lie on the diagonal axis. As visual aid, this diagonal is drawn in both scatter plots. Points depicted in the lower triangle of a scatter plot represent records for which the *f*_GI_ (*x*_1_*, x*_2_) model results in smaller mean squared error values and points in the upper triangle represent records where the *f*_mean_ (*x*_1_*, x*_2_) model performs better. There are a few outliers depicted in the lower triangle of the errors of the Mathews Griner data for records which yield an mean squared error value above 0.03with the *f*_mean_ (*x*_1_*, x*_2_) model. Investigating these records shows a huge difference in slope parameters of the conditional responses. To give an example, the slope parameters of the record that results in an mean squared error value above 0.04 for the *f*_mean_ (*x*_1_*, x*_2_) model are *s*_1_ = 1.2 and *s*_2_ = 9.6. The *f*_1*→*2_ (*x*_1_*, x*_2_) model gives an almost ten times higher error, which is caused by the surface being strongly contracted to the origin. The majority of errors scatter in the range of [0, 0.01] and are slightly above the diagonal. In the scatter plot on the left-hand side of Fig. 7, there is a clear tendency of records to scatter in the upper triangle, supporting the Wilcoxon signed-rank test result of the *f*_GI_ (*x*_1_*, x*_2_) model resulting in larger error values.

**Figure 7:**
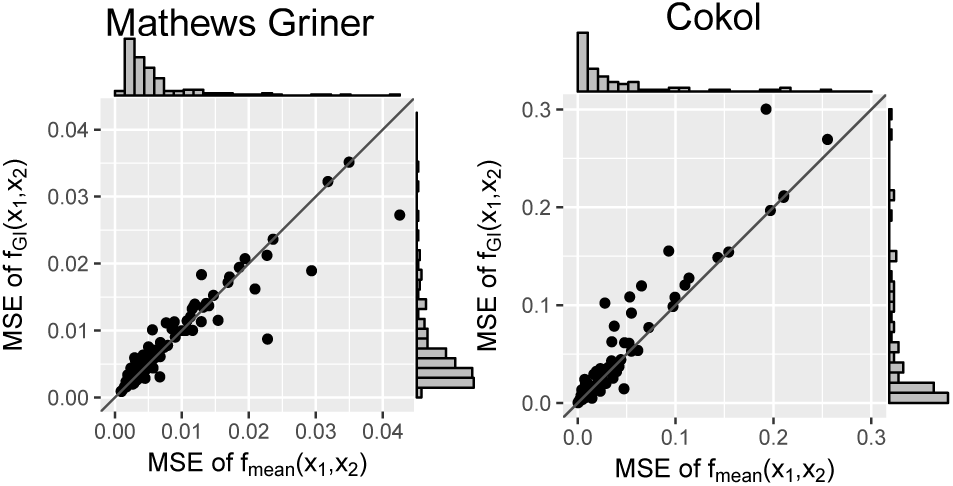
Mean squared error between the measured and the expected responses of the f_mean_ (x_1_, x_2_) and fGI (x_1_, x_2_) model. To better qualify the differences in mean squared error, the diagonal is depicted. The distribution of the models’ errors is given in histograms plotted on the axes.

To further investigate the difference between the implicit and explicit formulation derived from the Loewe Additivity principle, we conduct a small benchmarking test. We compare the computation time of the null reference models *f*_GI_ (*x*_1_*, x*_2_) (Eq. 5) and *f*_mean_ (*x*_1_*, x*_2_) (Eq. 14). For this, we use the data set from a study conducted by Yonetani and Theorell [19] which is believed to have no synergistic or antagonistic effect [20]. The data represents the inhibition of horse liver alcohol dehydrogenase by two inhibitors, ADP ribose and ADP. The data was used in an analysis of Chou and Talalay [9]. We fit the conditional parameters as described in Appendix E. For benchmarking, we use the microbenchmark package [21]. It runs each calculation per default 100 times. The median time to compute the explicit formulation of Loewe Additivity is 280 times faster than the implicit one (comparing to *f*_GI_ (*x*_1_*, x*_2_)) despite the double computing effort created by the Explicit Mean Equation model. Both, *f*_2*→*1_ (*x*_1_*, x*_2_*, y*) and *f* (*x, x, y*), have to be computed. Further results for the benchmark test on the null reference models are shown in Fig. 8 in Appendix F.

**Figure 8:**
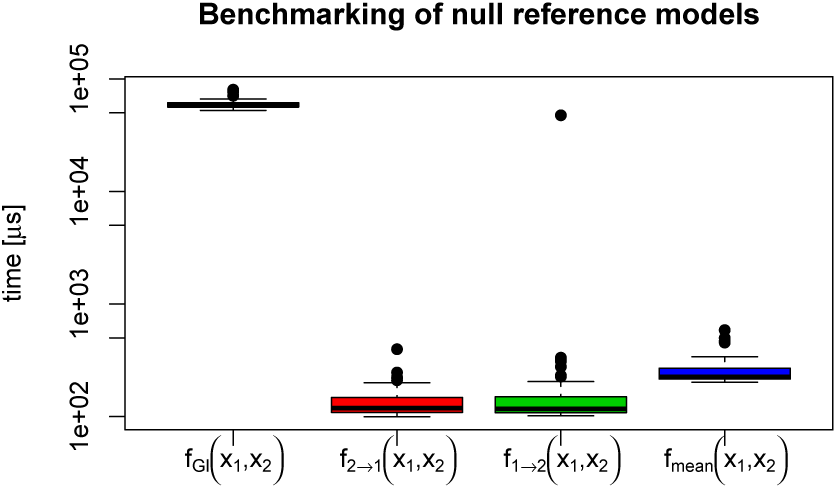
Benchmarking times on a log-scale for the four models fGI (x_1_, x_2_) (Eq. 5), the explicit formulation (Eq. 8 and Eq. 9), and its mean formulation fmean (x_1_, x_2_) (Eq. 14).

## VI. Discussion

With the rise of high-throughput methods, there is a huge opportunity to investigate compound combinations for synergistic effects. Especially with a first success in a synergy study in vivo mice [22], there is an urge to develop reliable methods to screen for promising combinations. Loewe Additivity is one of the most popular principles to investigate synergistic effects in compound combination studies. With the mathematical formulation in the first part of this study we are to our knowledge the first to have developed the theoretical background and the consistency condition of Loewe Additivity. Further, this mathematical derivation led to an explicit formulation of Loewe Additivity which underlines the arbitrariness of models derived from the Loewe Additivity principle. As commented upon before in [6, 13], we showed in two data sets that the LACC is often violated. These violations lead to differing predictions for different null reference models. Despite the common violation of the LACC, the *f*_GI_ (*x*_1_*, x*_2_) is popular. Therefore, it is important to tackle the biological question of which interaction to expect in case LACC is violated. We introduced the explicit model which is equivalent to the General Isobole Equation under the LACC and spans a similar surface if the LACC is violated. In two non-interactive high-throughput data sets we found our new Explicit Mean Equation null reference model to show smaller bias values than those of the General Isobole Equation model. This is a consequence of the more contracted surface to the origin if the LACC is violated. Further, we found the explicit Explicit Mean Equation to have smaller mean squared errors than the General Isobole Equation. These findings provide for an explicit model to replace the standard implicit model, both based on the Loewe Additivity principle. Additionally, the explicit model speeds up the computation time by a factor of roughly 250. In a large high-throughput experiment with 10, 000 response matrices this would reduce computing time from 20 hours to 5 minutes. We herewith provide a first step into the direction of improving the biological and numerical issues that follow from the Loewe Additivity principle.

## Acknowledgments

We thank Joris Mooij for his help in proving the LACC. We thank Bhagwan Yadav for the sharing of the code used for the analysis in [2] and Murat Cokol for the sharing of the data and analytical insights from [3].

## A. Loewe Additivity Consistency Condition

### Theorem 1.

*Proof.* For notational convenience we define 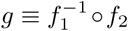 thus 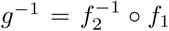 The function *g* maps dose *x*_2_ to its effect-equivalent dose *x*_1_. The LACC can now be written as:

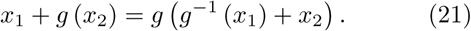

“⇒” We first provide a proof for the LACC to hold if the equivalent doses are proportional, meaning

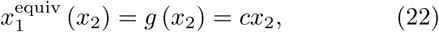

and thus

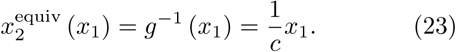

We therefore have, by rewriting Eq. 21,

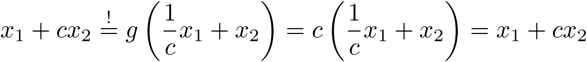

“⇐” Proving the theorem in the other direction, we assume that the LACC holds. Starting from the statement in Eq. 21 we define the function *h* (*x*_1_*, x*_2_) as the difference of these two equations:

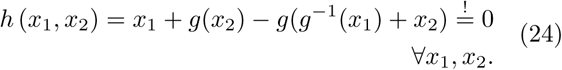

As *h* (*x*_1_*, x*_2_) is constant for all *x*_1_*, x*_2_, its first derivative has to be zero:

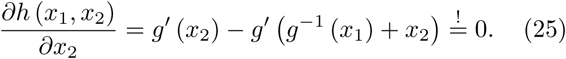

To further exclude that the inverse of *g* on *x*_1_ is always equal to zero, we take the derivative with respect to *x*_1_, which yields

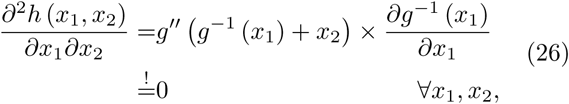

from which one can deduct that *g”( g*^*-*1^ (*x*_1_) + *x*_2_) has to be zero for all *x*_1_*, x*_2_. This implies that *g* (*x*_2_) is linear in *x*_2_. Thus:

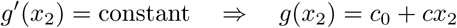

and since *g*(0) = 0, one obtains *c*_0_ = 0. Substituting this result in Eq. 6, one gets:

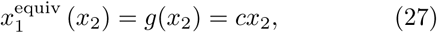

which shows equivalent doses to be proportional.

## B. General Isobole Equation under LACC

**Corollary 1.** *If the Loewe Additivity Consistency Condition in Eq. 10 holds,* (1) *f*_*GI*_ (*x*_1_*, x*_2_) = *f*_2*→*1_ (*x*_1_*, x*_2_) = *f*_1*→*2_ (*x*_1_*, x*_2_) *and* (2) *the isoboles are parallel.*

*Proof.* (1) Assume a response level denoted by *y*^*\**^, then the doses of compounds 1 and 2 that cause this effect by themselves are given by 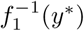 and 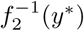 being related by concentration scaling, 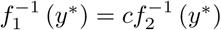 (Eq. 11 and 12). The isobole for effect *y*^*\**^ is given by

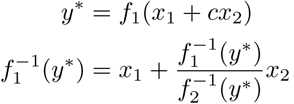

which leads to the linear isobole equation depicted in Eq. 5, assuming *f* is a continuous and monotonic response:

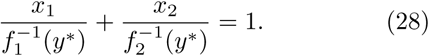

(2) For a given *y*^*\**^, this linear isobole equation gives the contour lines of the response surface. Since the two concentrations 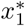 and 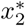 are related by linear scaling (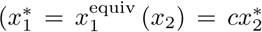, see Eq. 11), the isoboles for different effect levels *y*^*\**^ are parallel. This becomes clear by replacing 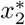 with its equivalent 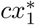 in the linear isobole equation and solving then for *x*_2_:

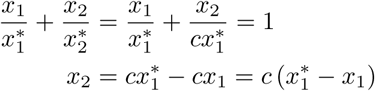

where 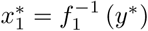 Therefore, *x*_2_ is linearly dependent with a fixed slope parameter *c >* 0. This results in an isobole with slope *c* for any effect *y*^*\**^ that is reached by 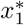, and therefore the isoboles are parallel.

## C. Consistency condition for Hill curve

**Corollary 3.** *If the* Loewe Additivity Consistency Condition *in Eq. 10 holds with f*_1_ *and f*_2_ *taking the form of two Hill curves, then the slopes and effect ranges y*_0_ *and y*_*∞*_ *of the Hill curves must be the same. Further, the proportionality factor c takes the form of a fraction of the EC50 value of the drug to be expressed in terms of the other divided by the EC50 value of this other drug, resulting in* 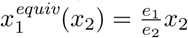

For the Loewe Additivity Consistency Condition,

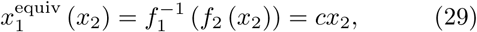

to be fulfilled when using the Hill curve for *f*_*j*_*, j*ϵ{1, 2 }, which is of the form

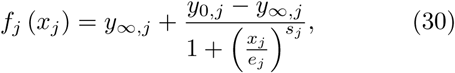

with its inverse being of the form

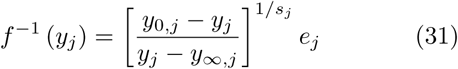

one gets:

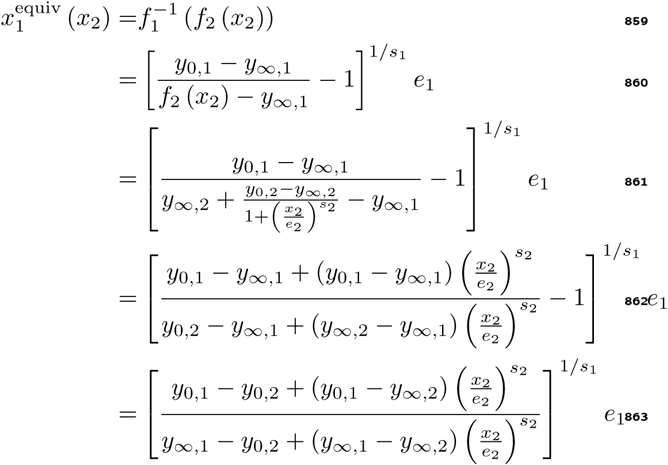

Hence, in order for 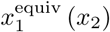 to be constant *y*_*∞,*1_ = *y*_*∞,*2_, which gives:

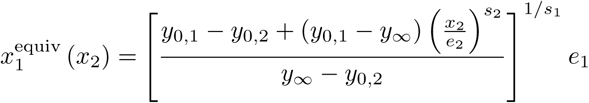

and *y*_0,1_ = *y*_0,2_, simplifying to:

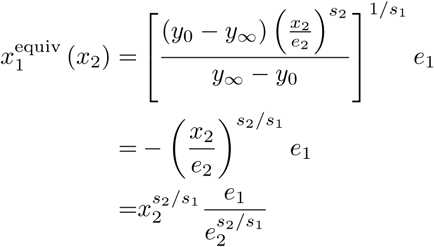

which is constant if *s*_2_*/s*_1_ =1*⇔s*_1_ = *s*_2_.

The Loewe Additivity Consistency Condition is therefore fulfilled for the Hill curves in the same ranges, *y*_*∞,*1_ = *y*_*∞,*2_ and *y*_0,1_ = *y*_0,2_ and with the same slope *s*_1_ = *s*_2_.

## D. Violation of the Loewe Additivity Consistency Condition

When the LACC applies, the General Isobole Equation, *f*_GI_ (*x*_1_*, x*_2_), and both explicit solutions, *f*_2*→*1_(*x*_1_*, x*_2_) and *f*_1*→*2_(*x*_1_*, x*_2_), are equivalent. When the LACC fails, one may wonder how to combine the two explicit solutions such that they are still close to the solution of the General Isobole Equation Eq. 5. Since the solution of the General Isobole Equation Eq. 5 is symmetric, it makes sense to take the (weighted) mean of the two solutions *f*_2*→*1_(*x*_1_*, x*_2_) and *f*_1*→*2_(*x*_1_*, x*_2_). Therefore, we define

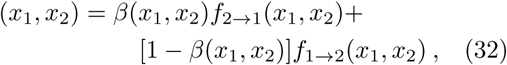

with *β*(*x*_1_*, x*_2_) a weighting function. We wonder how to choose *β*(*x*_1_*, x*_2_) such that *f*_mean_ (*x*_1_*, x*_2_) *≈ f*_GI_ (*x*_1_*, x*_2_) under mild violations of the LACC.

In case the LACC holds, we have

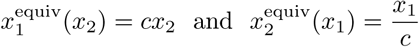

i.e. the equivalent doses are proportional to the original doses. To study mild violations of LACC, which we will refer to as LACC-*ϵ*, we add a small quadratic term, i.e., consider

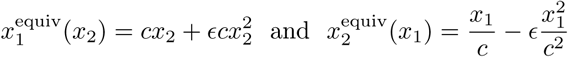

such that we have the consistency equations:

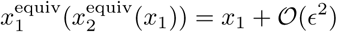

and

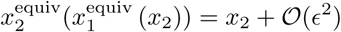

where we take ϵ to be small so that we can ignore all second and higher order terms in ϵNote the different dependence on *c* in the two quadratic terms. These are necessary for the consistency condition to hold. Our goal is now to find *β*(*x*_1_*, x*_2_) such that under LACCϵwe still have *f*_mean_ (*x*_1_*, x*_2_) = *f*_GI_ (*x*_1_*, x*_2_), or, to put it differently, such that *f*_mean_ (*x*_1_*, x*_2_) still satisfies the General Isobole Equation.

### Theorem 2.

*Under LACC*_*-*ϵ_*, the simple arithmetic mean f*_*mean*_ (*x*_1_*, x*_2_) *with β*(*x*_1_*, x*_2_) = 1*/*2 *satisfies the General Isobole Equation.*

*Proof.* As an intermediate step, we note that we can rewrite *f*_2_ into *f*_1_ and vice versa through 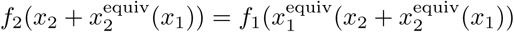 and 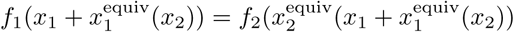
With

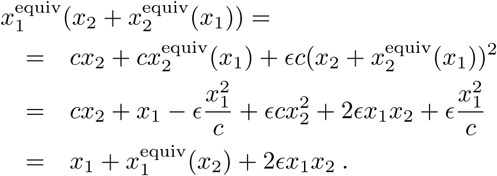

And

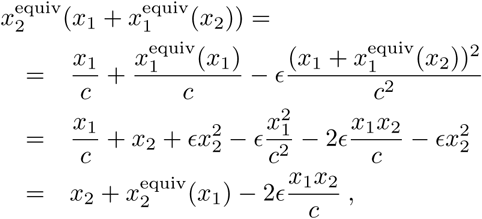

where, here and in the following, we ignore second and higher order terms inϵ.

For *f*_mean_ (*x*_1_*, x*_2_) to satisfy the General Isobole Equation, we need

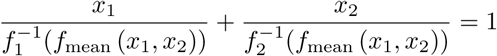

Luckily, using the above expressions, we have two ways to rewrite *f*_mean_ (*x*_1_*, x*_2_): in terms of *f*_1_,

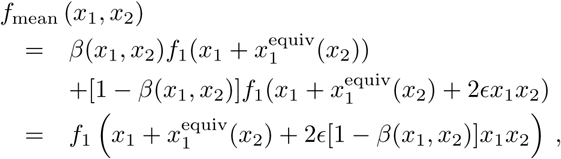

and in terms of *f*_2_,

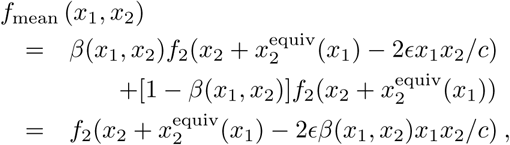

where the last step follows from the observation that, again keeping track of just the first order terms inϵ,

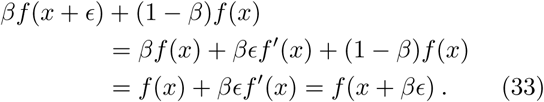

Plugging these two formulations at the obvious places in the General Isobole Equation, we obtain

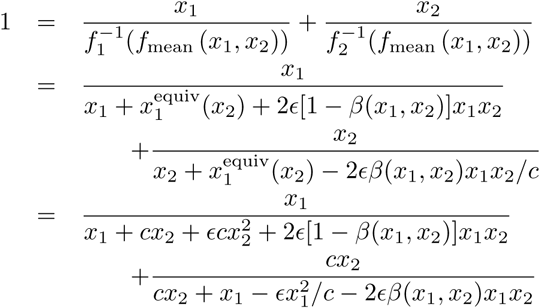

Further expansion in ϵyields

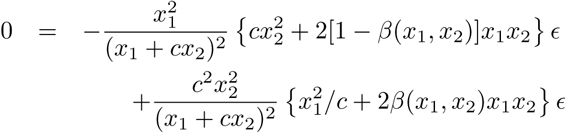

So, for the first order term in ϵto cancel, we must have

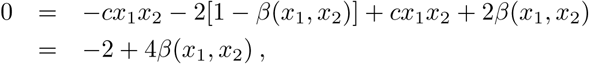

with solution *β*(*x*_1_*, x*_2_) = 1*/*2: simple equal weighting.

## E. Data Cleaning, Fitting of Hill Curve and Parameter Estimation for Implicit Models

A first step of processing the data includes an outlier analysis of the raw reads by fitting a spline surface and deleting outliers. Further, we explain how to calculate the implicit response values of the *f*_GI_ (*x*_1_*, x*_2_) model.

To detect outliers, we fit the data to a general additive model (GAM) using thin plate splines [23]. We use the method gam() of the mgcv-package [24]. Every data point is rejected for which its absolute residual value is larger than three times the inter-quantile range of all residuals of a given record. For the Mathews Griner data, this leads to 125 records out of the 466 where a mean of 1.59 outliers were excluded per record. A maximum of 6 outliers was detected once. Similarly, we excluded 4.21 data points for the Cokol data on 150 of the total 200 records with a maximum of 13 data points.

The two conditional responses of a record are fitted in parallel to two Hill functions in the form of Eq. 1 with the drc package [16]. Unlike other synergy analyses such as [2], the response at zero concentration *y*_0_ is not fixed but only constrained to be the same for both response curves. The other Hill parameters, *y*_*∞*_*, s* and *e* are fitted for both compounds individually.

The *f*_GI_ (*x*_1_*, x*_2_) model is an implicit model for the response *y*. Therefore, a root finder is used to find a response 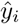 for a given parameter set Θ= {*y*_0_*, y*_*∞,j*_*, e*_*j*_*, s*_*j*_ *}*, and concentrations *x*_*i,*1_ *x*_*i,*2_. For finding such a root the standard implementation of a root finder in the R stats package, uniroot() [25], is used which uses the BrentDekker-van Wijngaarden algorithm [26, Chapter 9]. As convergence criterion we used 1.22 − 10^*-*4^. To ensure the existence of the inverse values of the Hill curve at a given response *y*, *f* ^*-*1^ (*y*), the responses are limited to the range for which the Hill curve is defined, [*y*_0_*, y*_*∞*_]. Responses outside this range are set to the closest range limits, i.e. if *y > y*_0_ *⇒ y* = *y*_0_ and if *y < y*_*∞*_ *⇒ y* = *y*_*∞*_.

## F. Benchmark Test on Generated Data from Yonetani

To show the speed advantage of the explicit formulation derived from the Loewe Additivity principle, we conducted a benchmarking test. We compare the computation time of the null reference models GI (Eq. 5), *f*_2*→*1_ (*x*_1_*, x*_2_) (Eq. 8), *f*_1*→*2_ (*x*_1_*, x*_2_) (Eq. 9), and Explicit Mean Equation (Eq. 14). For this, we use the data set from [19] which is known to be non-interactive [20]. We fit the conditional parameters as described in Appendix E. For the benchmarking test, we made use of the microbenchmark package [21]. It runs each calculation per default 100 times.

A visualization of the runtime is depicted in Fig. 8 The explicit formulations are clearly faster in computing time than the implicit ones.

## G. Geometric Mean Model

Analogously to the weighted mean, we take the geometric mean of the two explicit models *f*_2*→*1_ (*x*_1_*, x*_2_) and *f*_1*→*2_ (*x*_1_*, x*_2_) depicted in Eq. 8 and 9

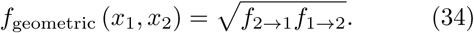

In parallel to the Explicit Mean Equation, the geometric mean is the least sensitive to a violation of the LACC with an equal weighting of both formulations *f*_2*→*1_ and *f*_1*→*2_: Assume the Explicit Geometric Mean Equation model with weights *β*(*x*_1_*, x*_2_):

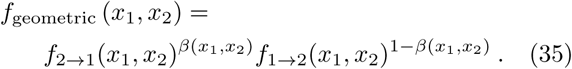

Obviously, under LACC we have *f*_geometric_ (*x*_1_*, x*_2_) = *f*_mean_ (*x*_1_*, x*_2_) = *f*_GI_ (*x*_1_*, x*_2_) for any choice of *β*(*x*_1_*, x*_2_).

### Theorem 3.

*Under LACC-*_ϵ,_ *the (weighted) geometric mean f*_*geometric*_ (*x*_1_*, x*_2_) *equals the arithmetic mean f*_*mean*_ (*x*_1_*, x*_2_) *with the same β*(*x*_1_*, x*_2_).

*Proof.* Following the exact same line of reasoning as in the proof of Theorem 4 up to Eq. 33 in Appendix D, again ignoring second and higher order terms inϵ, we get

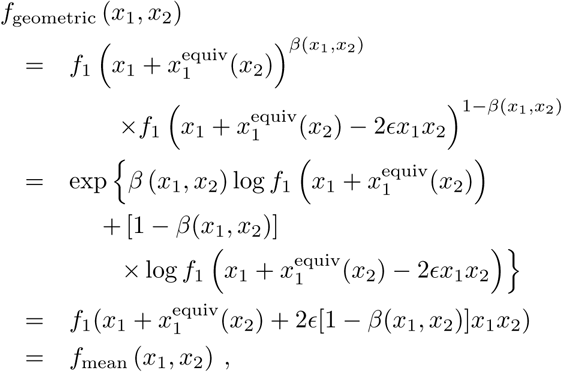

where we applied the reasoning of (33) to the term in the exponent.

This then gives the following corollary:

**Corollary 4.** *Under LACC-*_ϵ_*, the simple geometric mean f*_*geometric*_ (*x*_1_*, x*_2_) *with β*(*x*_1_*, x*_2_) = 1*/*2 *satisfies the General Isobole Equation*

Mathews GrinerCokol

Analogously to Fig. 5, the Explicit Geometric Mean Equation model takes different shapes for the different violations of LACC, as depicted in Fig. 3. They are depicted in Fig. 9

**Figure 9:**
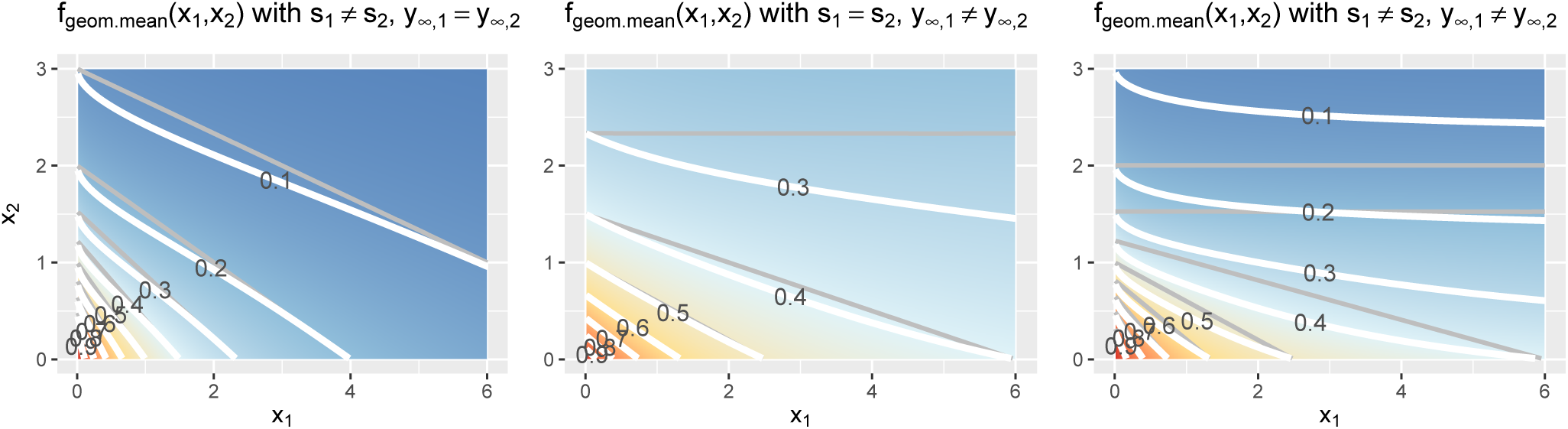
Contour lines of the fmean (x_1_, x_2_) model for three different scenarios, where the LACC is violated: from left to right: the slopes are different, s_1_ ≠ s_2_, here depicted with s_1_ = 1, s_2_ = 2, y_∞_ = 0, or the maximal effect values differ, y_∞_,_1_ = y_∞_,_2_, here shown with s = 1, y_∞_,_1_ = 0.3, y_∞_,_2_ = 0 or both are different, here shown with s_1_ = 1, s_2_ = 2, y_∞_,_1_ = 0.3, y_∞_,_2_ = 0. The remaining two parameters of the Hill curve are set equally for all figures to y_0_ = 1 and e = 1 are equal.

We compute the mean squared error values of each non-interactive record and compare them with the mean squared errors of the *f*_GI_ (*x*_1_*, x*_2_) model. The scatter plots for the two data sets are depicted in Fig. 10. A Wilcoxon signed-rank on both error value sets for the *f*_GI_ (*x*_1_*, x*_2_) and the *f*_geometric_ (*x*_1_*, x*_2_) model, with the alternative hypothesis being that the errors of *f*_GI_ (*x*_1_*, x*_2_) are larger, give the following *p*-values: for the Mathews Griner data set: 6.38 − 10^*-*6^ and for Cokol 7.26 − 10^*-*4^.

**Figure 10:**
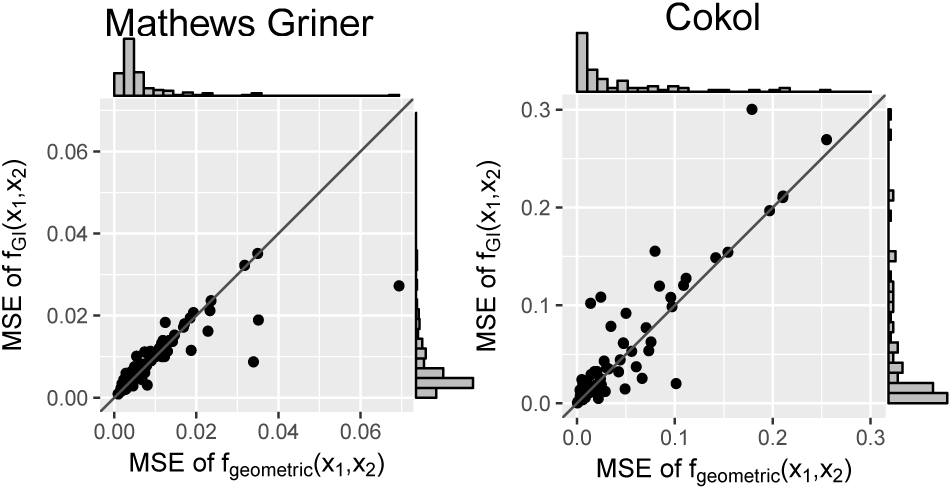
Mean squared error between the measured and the expected responses of the fgeometric (x_1_, x_2_) and fGI (x_1_, x_2_) model. To better qualify the differences in mean squared error, the diagonal is depicted. The distribution of the models’ mean squared error is given in histograms plotted on the axes.

## References

[1] S Loewe. Die quantitativen Probleme der Phar-makologie. Ergebnisse der Physiol., 27(1):47–187, 1928.

[2] Bhagwan Yadav, Krister Wennerberg, Tero Ait-tokallio, and Jing Tang. Searching for Drug Syn-ergy in Complex Dose-Response Landscapes Using an Interaction Potency Model. Comput. Struct. Biotechnol. J., 13:504–513, 2015.

[3] Murat Cokol, HN Chua, and Murat Tasan. Systematic exploration of synergistic drug pairs. Mol. Syst. Biol., 7(544), 2011.

[4] J Lehar, AS Krueger, W Avery, AM Heilbut, and LM Johansen. Synergistic drug combinations improve therapeutic selectivity. Nat Biotechnol, 27(7):659–66, 2009.

[5] William R. Greco, Gergory Bravo, and John C. Parsons. The Search for Synergy: A Critical Review from A Response Surface Perspective. Pharmacol. Rev., 47(2):331–385, 1995.

[6] Nori Geary. Understanding Synergy. Am. J. Phys-iol. Endocrinol. Metab., 304(3):E237–E253, 2012.

[7] Julie Foucquier and Mickael Guedj. Analysis of drug combinations: current methodological landscape. Pharmacol. Res. Perspect., 3(3), 2015.

[8] C I Bliss. The toxicity of poisons applied jointly. Ann. Appl. Biol., 26(3):585–615, 1939.

[9] Tin-Chao Chou and Paul Talalay. A Simple Generalized Equation for the Analysis of Multiple Inhibitions of Michaelis-Menten Kinetic Systems. J. Biol. Chem., 252(18):6438–6442, 1977.

[10] Giovanni Y. Di Veroli, Chiara Fornari, Dennis Wang, Severine Mollard, Jo L Bramhall, Frances M Richards, and Duncan I Jodrell. Combenefit: An in-teractive platform for the analysis and visualisation of drug combinations. Bioinformatics, 32(18):2866–2868, 2016.

[11] Joseph Lehar, Grant R Zimmermann, Andrew S Krueger, Raymond a Molnar, Jebediah T Ledell, Adrian M Heilbut, Glenn F Short, Leanne C Giusti, Garry P Nolan, Omar a Magid, Margaret S Lee, Alexis a Borisy, Brent R Stockwell, and Curtis T Keith. Chemical combination effects predict con-nectivity in biological systems. Mol. Syst. Biol., 3(80):80, 2007.

[12] Desiree Y Baeder, Guozhi Yu, Nathanael Hoze, Jens Rolff, and Roland R Regoes. Antimicrobial combi-nations: Bliss independence and Loewe additivity derived from mechanistic multi-hit models. Philos. Trans. B, 371, 2016.

[13] Ronald J Tallarida. Revisiting the Isobole and Related Quantitative Methods for Assessing Drug Synergism. J. Pharmacol. Exp. Ther., 342(1):2–8, 2012.

[14] A. V. Hill. The possible effects of the aggregation of the molecule of hemoglobin on its dissociation curves. J. Physiol., 40:iv–vii, 1910.

[15] Sylvain Goutelle, Michel Maurin, Florent Rougier, Xavier Barbaut, Laurent Bourguignon, Michel Ducher, and Pascal Maire. The Hill equation: A review of its capabilities in pharmacological modelling. Fundam. Clin. Pharmacol., 22(6):633–648, 2008.

[16] Christian Ritz and Jens C Strebig. drc: Analysis of Dose-Response Curves, 2016.

[17] M C Berenbaum. What is Synergy? Pharmacol. Rev., 2(41):93–141, 1989.

[18] Lesley A Mathews Griner, Rajarshi Guha, Paul Shinn, Ryan M Young, Jonathan M Keller, Dongbo Liu, Ian S Goldlust, Adam Yasgar, Crystal McK-night, Matthew B Boxer, Damien Y Duveau, Jian-Kang Jiang, Sam Michael, Tim Mierzwa, Wenwei Huang, Martin J Walsh, Bryan T Mott, Paresma Patel, William Leister, David J Maloney, Christo-pher A Leclair, Ganesha Rai, Ajit Jadhav, Brian D Peyser, Christopher P Austin, Scott E Martin, An-ton Simeonov, Marc Ferrer, Louis M Staudt, and Craig J Thomas. High-throughput combinatorial screening identifies drugs that cooperate with ibru-tinib to kill activated B-cell-like diffuse large B-cell lymphoma cells. Proc. Natl. Acad. Sci. U. S. A., 111(6):2349–54, 2014.

[19] T. Yonetani and H. Theorell. Studies on liver al-cohol dehydrogenase complexes: III. Multiple inhi-bition kinetics in the presence of two competitive inhibitors. Arch. Biochem. Biophys., 106:243–251, 1964.

[20] Ting-Chao Chou and Paul Talalay. Quantitative analysis of dose-effect relationships: the combined effects of multiple drugs or enzyme inhibitors. Adv. Enzyme Regul., 22:27–55, 1984.

[21] Olaf Mersmann. microbenchmark: Accurate Timing Functions, 2015.

[22] Barbara M Grüner, Christopher J Schulze, Dian Yang, Daisuke Ogasawara, Melissa M Dix, Zoë NRogers, Chen-Hua Chuang, Christopher D McFar-land, Shin-Heng Chiou, J Mark Brown, Benjamin F Cravatt, Matthew Bogyo, and Monte M Winslow. An in vivo multiplexed small-molecule screening platform. Nat. Methods, 13(September), 2016.

[23] Simon Wood. Generalized Additive Models: an introduction with R. Biometrics, 62(4):392, 2006.

[24] Simon Wood. mgcv: Mixed GAM Computation Vehicle with GCV/AIC/REML Smoothness Estima-tion, 2016.

[25] R Core Team. R: A Language and Environment for Statistical Computing. R Foundation for Statistical Computing, Vienna, Austria, 2016.

[26] William H Press, Saul a Teukolsky, William T Vet-terling, and Brian P Flannery. Numerical Recipes: The Art of Scientific Computing, volume 1. Cambridge University Press, Cambridge, 3 edition, 2007.

